# Finding Overlapping Rmaps via Gaussian Mixture Model Clustering

**DOI:** 10.1101/2021.07.16.452722

**Authors:** Kingshuk Mukherjee, Massimiliano Rossi, Daniel Dole-Muinos, Ayomide Ajayi, Mattia Prosperi, Christina Boucher

## Abstract

Optical mapping is a method for creating high resolution restriction maps of an entire genome. Optical mapping has been largely automated, and first produces single molecule restriction maps, called Rmaps, which are assembled to generate genome wide optical maps. Since the location and orientation of each Rmap is unknown, the first problem in the analysis of this data is finding related Rmaps, i.e., pairs of Rmaps that share the same orientation and have significant overlap in their genomic location. Although heuristics for identifying related Rmaps exist, they all require quantization of the data which leads to a loss in the precision. In this paper, we propose a Gaussian mixture modelling clustering based method, which we refer to as OMclust, that finds overlapping Rmaps without quantization. Using both simulated and real datasets, we show that OMclust substantially improves the precision (from 48.3% to 73.3%) over the state-of-the art methods while also reducing CPU time and memory consumption. Further, we integrated OMclust into the error correction methods (Elmeri and cOMet) to demonstrate the increase in the performance of these methods. When OMclust was combined with cOMet to error correct Rmap data generated from human DNA, it was able to error correct close to 3x more Rmaps, and reduced the CPU time by more than 35x. Our software is written in C++ and is publicly available under GNU General Public License at https://github.com/kingufl/OMclust

## 1 INTRODUCTION

A restriction map is defined as a map that records the locations of one or more specific short nucleotide sequences called restriction sites, across a DNA sequence. Restriction maps are generated by treating DNA molecules with special enzymes, known as restriction enzymes, that recognizes the restriction sites in the DNA sequence and cleaves the DNA wherever these sites occur. Optical mapping is a sequencing technique that produces high resolution restriction maps of an entire genome, giving it a unique numeric representation. It is used alone or in concert with sequence data, i.e., assembling long and short reads [1, 8, 12, 27], scaffolding assembled regions [4, 20, 28], detecting misassemblies in draft genomes [17, 21] and finding structural variations [5, 11].

The main laboratory steps of optical mapping are as follows. First, the DNA of multiple cells of the same organism are untangled and randomly sheared to produce a large collection of DNA molecules which are then stretched and held in place on a slide and observed under a fluorescent microscope. Next, a restriction enzyme is selected and applied to the DNA which cleaves them at specific sites, called cut sites. The cleaved DNA fragments are size estimated and ordered, resulting in a single molecule restriction map for each sheared DNA molecule. These single molecule restriction maps called *Rmaps* are the raw data. For example, digesting a DNA sequence with the restriction enzyme BspQI which recognizes the site GCTCTTC, we may obtain the following Rmap *R* = [1213, 7129, 19 632, 2845, 11 754, 1935, 9775, 4005, 3854, 17 432] .From this we can deduce that the first occurrence of GCTCTTC in the DNA sequence is at location 1213 base-pairs(bp) followed by 8,342 bp and so on. A large portion of this process has been automated, and with the automation the resulting data has become increasingly more accurate over the past decade. For example, BioNano’s nanochannel has significantly improved the accuracy and accessibility of the data [7].

After the Rmaps are produced they are assembled to produce genome wide optical maps, which are synonymous to assembled contigs from sequence data, i.e., they usually do not span an entire chromosome but span significantly larger genomic regions than the raw data. Also, similar to a sequence read, the location and orientation of an individual Rmap in the genome is unknown. For example the two Rmaps, *R*_1_ = [1213, 7129, 19 632, 2845, 11 754, 1935] and *R*_2_ = [2845, 11 754, 1935, 9775, 4005, 3854, 17 432] can be assembled to produce the consensus optical map *R* = [1213, 7129, 19 632, 2845, 11 754, 1935, 9775, 4005, 3854, 17 432]. In a genome wide optical map, we see that *R*_1_ should occur and then be followed by *R*_2_. In addition, we note that uncorrected Rmaps contain the following types of errors: (a) *sizing error*, meaning that the estimation of the size of a fragment in the Rmap differs from the true size; (b) *added cut sites*, meaning a cut site occurs accidentally and results from the splicing of a single fragment into two or more smaller fragments; and lastly (c) *deleted cut sites*, meaning a cut site should have occurred but was missed and two or more smaller fragments are merged into a single one. Using our example above, a realistic representation of the two Rmaps with errors is as follows: 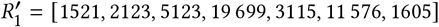 and 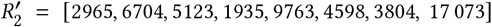 where missed cut sites are introduced in the second fragment of *R*_1_ and second fragment of *R*_2_, and sizing errors are introduced to all fragments from both Rmaps.

The first step in assembling Rmaps is to find all pairwise alignments, which are then used to error correct the Rmaps, and finding overlaps between the error corrected data. However, due to the error profile of Rmap data, finding pairwise alignments between Rmaps is challenging. Dynamic programming remains the only robust method for finding pairwise alignments between Rmaps as existing alignment methods largely focus on finding alignments between assembled genome wide optical maps [13, 15]. Unfortunately, dynamic programming is computationally expensive and is unable to scale to even moderately large sized genomes, such as the human genome [18]. Therefore, all existing error correction methods (cOMet and Elmeri) use heuristics to filter out pairs of Rmaps that are likely to not have significant pairwise alignment, and then find alignments between the remaining pairs [19, 22]. These methods, as well as other optical mapping analysis methods, require the data to be first quantized, which involves assigning all fragment sizes into discrete bins, and replacing all fragment sizes in the same bin with a pre-defined size. For example, if any fragment size within the range [1500, 2000] are binned together and assigned the value 1750, then 1625, 1855, and 1999 would be replaced with 1750 in their corresponding Rmaps. While these heuristics for finding pairs of related Rmaps are efficient, the accuracy of these heuristics has not been fully explored. Ideally, any method that filters for pairwise alignments should have high precision since the objective is to maximize the fraction of the true pairwise alignments.

In this paper, we propose an efficient clustering-based method for finding related Rmaps with high precision, which does not require any quantization or noise reduction. We refer to our method as OMclust. It first performs a grid search to find the best parameters of the clustering model and replaces quantization by identifying a set of cluster centers and uses the variance of the cluster centers to account for the noise. Next, for finding related Rmaps, a cluster is assigned to each *k*-mer (i.e. *k* consecutive fragments of an Rmap) extracted from the Rmaps based on its proximity to the identified cluster centers. Finally, we call a pair of Rmaps as related if a number of their *k*-mers are assigned to the same cluster.

We implemented OMclust and compared it to the heuristics for finding related Rmaps in Elmeri [22] and cOMet [19]. OMclust achieved the highest precision, and was most efficient with respect to both time and memory. In particular on a simulated *E. coli* dataset, we show OMclust found the relations with a precision of 73.3%; whereas, Elmeri achieves 48.3% precision and cOMet achieve a precision of 5.1%. Next, using a dataset consisting of over 14 million Rmaps generated from human DNA, we evaluated the increase to the performance of cOMet using OMclust to find the set of related Rmaps. We demonstrated that the combination of OMclust and cOMet outperformed cOMet in its default setting; the combined method corrected close to three times more Rmaps (from 987,985 to 2,757,266), and reduced the CPU time by more than 35 times (from 23,637 to 638 CPU hours). Lastly, we note that Elmeri ran out of memory (exceeding 800 GB) on the human dataset so we were unable to perform the same comparison.

## 2 RELATED WORK

As previously mentioned, assembly and error correction require finding pairwise alignments – and hence, pairs of related Rmaps – between Rmaps. In this section, we will briefly describe error correction and assembly methods.

Two error correction methods are available for Rmaps: cOMet [19] and Elmeri [22]. Both methods use an heuristic to quantize the data, find all pairs of related Rmaps, align each Rmap to all of its related Rmaps, use this alignment to find a consensus, and error correct each Rmap by making it consistent to the consensus. In both methods, the quantization scheme replaces all fragments by discrete number bins that are found by dividing the fragment sizes by an integer *b* and then rounding off the quotient to the closest integer value. All sets of *k* consecutive Rmap fragments, called *k*-mers, are then extracted from the quantized Rmaps for some value of *k*. Using the example above, quantizing 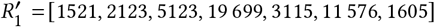 using a bucket size, *b* = 2000 transforms it into 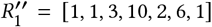. cOMet considers two Rmaps to be related if there exists at least *m* common *k*-mers, where *m* is an input parameter. After finding related Rmaps, cOMet finds all pairwise alignments using the dynamic programming alignment algorithm of Valouev *et al*. [25]. Although cOMet is shown to reliably reduce errors in Rmaps, it requires a considerable amount of time to find all pairwise alignments since the set of related Rmaps has low precision, i.e., considers a large number of Rmaps to be related. Elmeri improves upon the precision and recall of finding related Rmaps compared to cOMet [22] but it requires more than 10 times as much memory [22]. Elmeri replaces *k*-mers with (*k, ℓ*)-mers, which are defined as Rmap segments that add up to *ℓ* base pairs and contains at least *k* fragments. If a segment does not include at least *k* fragments then it is extended to the right until it covers *k* fragments. The (*k, ℓ*) -mers are extracted from quantized Rmaps, similar to cOMet, and a pair of Rmaps are deemed to be related if they have at least *m* common (*k, ℓ*) -mers. The related Rmaps are then aligned using a heuristic and Rmaps are error corrected based on the consensus of these alignments. Both Elmeri and cOMet are used in the evaluation of our method.

The traditional and more deterministic way of finding related Rmaps is to determine the alignment between all pairs of Rmaps. The alignment algorithm of Valouev *et al*. [25], Maligner [13], OMBlast [9] and Kohdista [16] are capable of finding pairwise alignments between Rmaps. The dynamic programming algorithm of Valouev *et al*. [25] can be seen as a modification of Smith-Waterman algorithm [24] which optimizes an alignment score that is referred to as *S-score*. The S-score is based on a probabilistic model that assumes that the fragment sizes follow an exponential distribution, the occurrence of a restriction site is an independent Bernoulli process, the number of false cuts in a genomic interval is a Poisson process, and the sizing error follows a gaussian distribution with mean zero and variance being a linear function of the true fragment size. Valouev *et al*. use the an S-score cutoff of 25 to distinguish between related and unrelated Rmaps; where two Rmaps whose S-score is greater than or equal to 25 are deemed to be related. We use the S-score later in the paper to evaluate the quality of relation calls. Although this method for alignment is robust to errors, it is impractical for even reasonably large datasets. OMBlast [9] uses a seed-and-extend method for finding alignments for Rmaps against a reference genome optical map by first building an index from the reference optical map. From this index, seeds are identified for a query Rmap and alignment is performed matching the segments between two consecutive seeds. Although this workflow is not primarily designed for pairwise alignment of Rmaps, it was shown to be able to generalize to that problem by considering each Rmap as a reference map and adjusting the parameters [16]. The index-based aligner Kohdista formulates the alignment problem as an automaton path-matching problem which builds an automaton from all the Rmaps and uses that for finding all alignments for Rmaps.

Since pairwise alignment of Rmaps is a critical step for Rmap assembly, the lack of an efficient solution to this problem has inhibited the development of Rmap assemblers. Currently there exists only two non-proprietary methods for Rmap assembly: the assembly method of Valouev *et al*. [26], and a de Bruijn graph-based assembler called rmapper [18]. The former uses the alignment method of Valouev *et al*. to first compute all pairwise alignments and uses a threshold alignment score to identify reliable overlaps among Rmaps. Then, it constructs an overlap graph and traverses it to construct genome wide optical maps. Due to the inefficiency of the pairwise alignment, this overlap-layout-consensus based method is only able to assemble small genomes with low coverage of data. rmapper is a recent assembly method that circumvents the problem of pairwise alignment of Rmaps by constructing a de Bruijn graph from all unique bi-labels extracted from Rmaps, where a bi-label is a pair of *k*-mers which are separated by a given length in base-pairs. This de Bruijn graph is then traversed to construct a genome wide optical map.

## 3 METHODS

### 3.1 Overview of OMclust

Our method requires an initial preprocessing phase, where, for a fixed *k*, we extract *k*-mers from all Rmaps in the training dataset. Then we train a Gaussian mixture model for finding *k*-mer cluster centers that best represents the dataset. Once the training phase is concluded, we find related Rmaps by extracting all *k*-mers from the test dataset and assigning each *k*-mer to a cluster center with a probability proportional to the proximity of the *k*-mer to the center of the cluster. Finally, we call two Rmaps as related if at least *m* of their *k*-mers are clustered together.

### 3.2 Gaussian mixture model clustering of *k*-mers

For an integer *k*, we first extract all *k*-mers from every Rmap in the training dataset. Here a *k*-mer refers to *k* successive fragments of the Rmap. If an Rmap has *n* fragments then a total of *n*− *k* +1 number of *k*-mers can be extracted from the Rmap. For example, say an Rmap has the following fragments: *R* = [1213, 7129, 19 632, 2845, 11 754, 1935]. Then the following 4-mers can be extracted: (1213, 7129, 19 632, 2845), (7129, 19 632, 2845, 11 754) and (19 632, 2845, 11 754, 1935). The extracted *k*-mers are stored in memory as a two-dimensional array of integers of size *N* × *k* where *N* is the number of *k*-mers.

The full parameter set can be compactly stated as 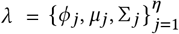. Next, we cluster the *k*-mers using multivariate Gaussian mixture model clustering [2]. This clustering approach assumes that in a sample *X* = *X*_1_, *X*_2_, …, *X*_*n*_ of observations from independent and identically distributed (i.i.d.) random variables, each *X*_*i*_ is a *k*-dimensional variable that comes from a finite-mixture, Gaussian-based probability density function made by *η* components, of the form:

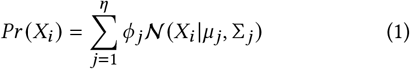

where *ϕ* _*j*_ is the prior for component *j* with constraints 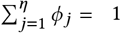 and *ϕ*_*j*_ ≥ 0 and 𝒩 (*X* |*μ*, Σ) is a *k*-dimensional Gaussian density function with mean *μ* and covariance matrix Σ defined as follows.

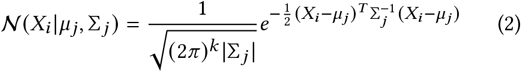

where |Σ_*j*_ | and 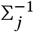 denote the determinant and inverse of Σ respectively and (*X*_*i*_ − *μ* _*j*_)^*T*^ denotes the transpose of (*X*_*i*_ − *μ* _*j*_). The most general form for a cluster is ellipsoidal, centered in *μ* _*j*_, while the co-variance matrix Σ_*j*_ determines other geometric features like orientation, volume, and shape. The simplest form for Σ_*j*_ is *λI*, which means a spherical Gaussian cluster, and requires only an additional parameter that is the standard deviation.

For a given number of clusters *η*, the mixture model’s parameters can be estimated via expectation-maximization [3]. However, determining the optimal number of clusters is not straightforward [6]. Standard approaches for this is to compute statistics such as the Bayesian Information Criterion (BIC) and the silhouette score. The BIC statistic calculates the in-sample model log-likelihood 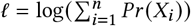 and penalizes it by the number of parameters *η* in relation to the sample size *n*, according to the formula BIC=*η*log(*n*) − 2*ℓ*. For the silhouette score, given a point, we compute the mean distance of the point from (i) all other points in its own cluster and from (ii) all points in next closest cluster. The silhouette score for the point is defined as the difference between (ii) and (i) divided by the greater of the two values. We use the mean of the silhouette score over all points to estimate the quality of the clustering. We also compute a similar statistic by recording the percentage of *k*-mers producing negative silhouette scores, where a lower value indicates a better clustering.

Hence, we fit a Gaussian mixture models and observed the variation of the three aforementioned statistic when the number of clusters are varied. In order to ameliorate possible variance in the statistics resulting from a single point estimate, we run the GMM optimization thrice for each cluster setting. Of note, we transformed the original values using the inverse hyperbolic sine function, `asinh(·)`, to reduce the value distribution skewness.

The clustering procedures were run using the Armadillo C++ library [23] which provides the class called *gmm*_*diag* for building and evaluating multi-threaded Gaussian mixture models. This class assumes spherical cluster shapes and uses a diagonal co-variance matrix.

The training of the clusters is the most expensive step in our algorithm. However, once we have trained clusters for a genome, these clusters can be used for classifying new data belonging to individuals of the same species, provided they are generated on the same platform (i.e. they will have same error rates) and use the same restriction enzyme. This is significant since optical maps are now mass-produced for human individuals and analyzed for structural variants and other markers.

### 3.3 Finding related Rmaps

Once we have fit the GMM, obtained the training data clusters and stored the clusters’ centers, we proceed to find related Rmaps from the test dataset as follows. First, we extract all *k*-mers from the Rmaps and for each *k*-mer, we store its corresponding Rmap in an array, which provides a mapping between *k*-mers and Rmaps. Next, we assign each *k*-mer to a single cluster which is defined by its nearest cluster center. For efficient assignment, we use a k-d tree to store all cluster centers and query the k-d tree repeatedly to find the nearest neighbor for each *k*-mer. We store the assignments in an array *A* of length equal to the number of *k*-mers in the test data where *A* [*i*] holds the cluster to which the *i*-th *k*-mer is assigned. Also, for every cluster *j*, we maintain a list *B* _*j*_ that stores the indices of the *k*-mers that are assigned to that cluster. For example, if *A* [*i*] = *j* then *B* _*j*_ will contain *i*.

For finding relations for an Rmap *R*_*i*_, we declare an array *C*_*i*_ of size equal to the number of Rmaps and initialize *C*_*i*_ with zeroes. Next, we traverse through the list of *k*-mers in *R*_*i*_, and for each *k*-mer, we visit the other *k*-mers that it is clustered together with using the data-structures *A* and *B*. For every such co-clustered *k*-mer, let *R*_*j*_ be the Rmap that it is extracted from, we note *R*_*j*_ by incrementing *C*_*i*_ [*j*] by 1. This implies that *R*_*i*_ and *R*_*j*_ has a pair of *k*-mers that are clustered together. We repeat this for all *k*-mers of *R*_*i*_. Finally, we traverse through *C*_*i*_ to identify the Rmaps that has *m* or more of their *k*-mers co-clustered with *R*_*i*_ and this set of Rmaps are deemed to be related to *R*_*i*_. We repeat this for all Rmaps in the dataset and report all pairs of Rmaps that have at least *m k*-mers in common.

## 4 EXPERIMENTS

In this section, we investigate the accuracy of our method OMclust and compare it with the method for finding related Rmaps used in cOMet and Elmeri. We performed all experiments on Intel E5-2698v3 processors with 100 GB of RAM running 64-bit Linux. For training, OMclust is run as a multi-threaded process on 64 CPUs in parallel. We used the Armadillo C++ linear algebra library which provides a fast and robust implementation of GMMs by employing multithreaded versions of the Expectation Maximisation (EM) and K-means algorithms reformulated into a Hadoop MapReduce-like framework. For finding related Rmaps, all methods are run on a single CPU, where not otherwise specified.

### 4.1 Metrics

We define two Rmaps as *related* if they are oriented in the same direction (i.e. both forward or both reverse with respect to the reference genome) and they overlap by at least 100,000 bp and 7 fragments. We chose this criteria to be consistent with prior work [22]. We refer to the set of all related Rmaps as the Positive (*P*) set. In addition, we refer to an Rmap relation as a True Positive (*T P*) if it is predicted by the method, and it is in *P*, and we refer to an Rmap relation as a False Positive (*F P*) if it is predicted by the method but it is not in *P*. Therefore, we define *precision* of a method as *T P*/ (*T P*+*F P*). The precision represents the percentage of correct Rmap relations a method made, and is also commonly referred to as the positive predictive value. Similarly, we define *recall* as*T P*/ *P*. The recall is also commonly referred to as sensitivity or true positive rate. We then consider *F*_*β*_ *score* which is the harmonic mean of the precision and recall where the recall is considered *β* times as important as the precision. We use *β* = 0.5. Therefore the *F*_*β*_ score is as follows (1+ *β*^2^) · *precision · recall* /(*β*^2^ · *precision* +*recall*). We are able to compute precision and recall only on simulated dataset.

Additionally, for a pair of related Rmaps, we compute the alignment score (called S-score) using the dynamic programming method of Valouev *et al*. [25]. For this alignment, we use the pairwise alignment module of the method which considers Rmap pairs that score above 25 as overlapping [26].

We further investigate the impact of using related Rmaps found by OMclust on the error correction modules of cOMet and Elmeri by replacing the related Rmaps found by their heuristics for finding related Rmaps with OMclust. Then we align the Rmaps before and after error correction to the reference optical map using the fit alignment module of the method of Valouev *et al*. and report the mean S-score of the alignment before and after error correction as well as the number of Rmaps whose S-score improved after error correction. These metrics are standard for evaluating efficacy of error correction [19, 22].

### 4.2 Datasets

We evaluated our method on a simulated *E. coli* bacterial dataset and a real human dataset. For our first experiment, we simulated two different Rmap datasets from *E. coli* K-12 substr MG1655 genome using the Rmap simulation software OMSim [14]. We used one of these datasets for training the clusters whereas the second is used for testing. We used the same simulation parameters and the same restriction enzyme, BspQI, for generating both datasets. We used the default error rate of OMSim for BspQI, which is a 15% rate of deleted cut sites, and 1 added cut site per 100 kbp. Since OMSim does not output the locations and orientations of the simulated Rmaps in the genome, we modified the software in order to record this information and used it to generate the ground truth for Rmap relations. The updated software is available at: https://github.com/asimas12/OMSIM_MODIFIED/tree/omsim_modified. The training dataset contains 8,095 Rmaps whereas the test dataset contains 8,051 Rmaps, both at 300x coverage.

For our second experiment, we used a real human Rmap dataset (Accession: SAMN01091029) generated for finding large structural variations from Levy-Sakin *et al*. [10]. This dataset consists of 14,692,103 Rmaps; which is 140x coverage. For training clusters for this dataset, we used OMSim to simulate human Rmap data from the human reference genome GRCh38 (accession number GCF_000001405.26) using the same restriction enzyme as was used to generate the real data. The simulated dataset contains 149,279 Rmaps; which is 25x coverage.

### 4.3 Results on simulated E. coli data

In order to estimate the optimal number of clusters for *E. coli*, we varied the number of clusters between 300 to 6,000 and plotting the BIC statistics, the mean Silhouette statistic and the percentage of negative Silhouette for various values of *k* (i.e., *k* = 4, *k* = 5, and *k* = 6) using the simulated training data. The plots are shown in Figures 1, 2, and 3. The BIC curve is maximized at around 3,800 clusters when *k* is equal to 4; while the mean silhouette curve is maximized at 2,200 clusters. The percent of negative silhouette scores hits a minimum at around 800 clusters. Since we have three different inflection points, we take the median of the three figures, which is 2,200, and set it as the optimal number of clusters for *k* = 4 for this dataset. Similarly, we find the inflection points to be at 3,800, 2,600 and 600 when *k* is equal to 5. Hence, we choose the median value 2,600 as optimal for *k* = 5. When *k* is equal to 6, we find inflection points at 4.200 in the BIC curve, and 800 in the percent of negative silhouette scores curve but do not find one in the mean silhouette curve. Hence, we choose the median of the two, i.e., 3,000 clusters was optimal for *k* = 6. We store these cluster centers from the training step to be used in testing. When *k* is equal to 4, the wall time for training is 2 minutes and 43 seconds using 64 CPUs. The corresponding CPU time was 2 hours 20 minutes and 43 seconds. When *k* is equal to 5, the wall time for training was 3 minutes and 1 seconds, and the CPU time was 2 hours 9 minutes and 3 seconds. Lastly, when *k* is equal to 6, the wall time for training was 3 minutes and 8 seconds, and the CPU time was 2 hours 23 minutes and 31 seconds.

**Figure 1:**
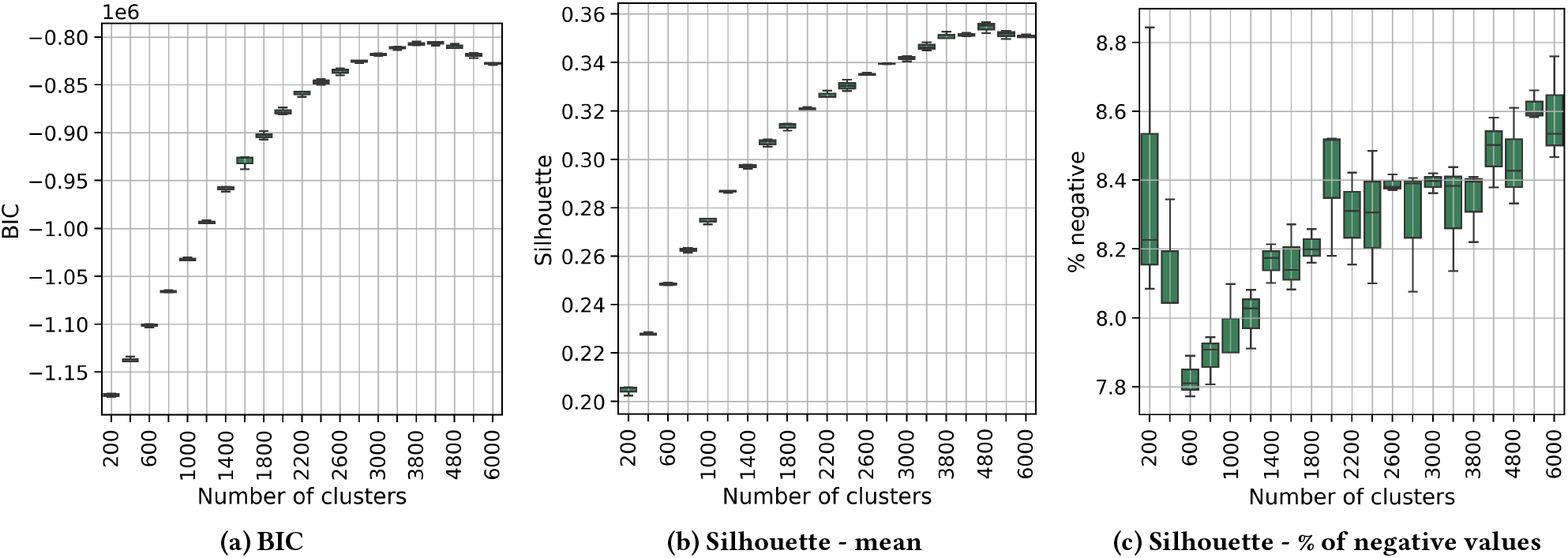
**Variation of BIC, mean silhouette scores and percentage of negative silhouette scores with respect to the number of clusters for *E. coli* for *k* = 4.**

**Figure 2:**
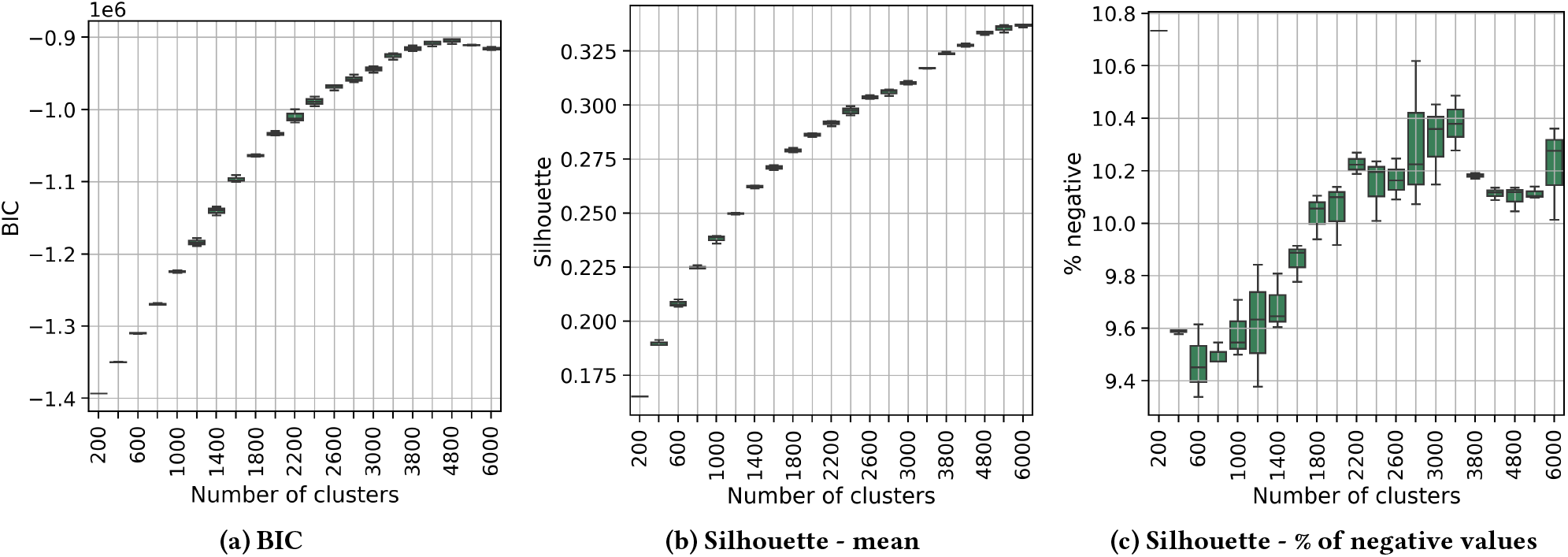
**Variation of BIC, mean silhouette scores and percentage of negative silhouette scores with respect to the number of clusters for *E. coli* for *k* = 5.**

**Figure 3:**
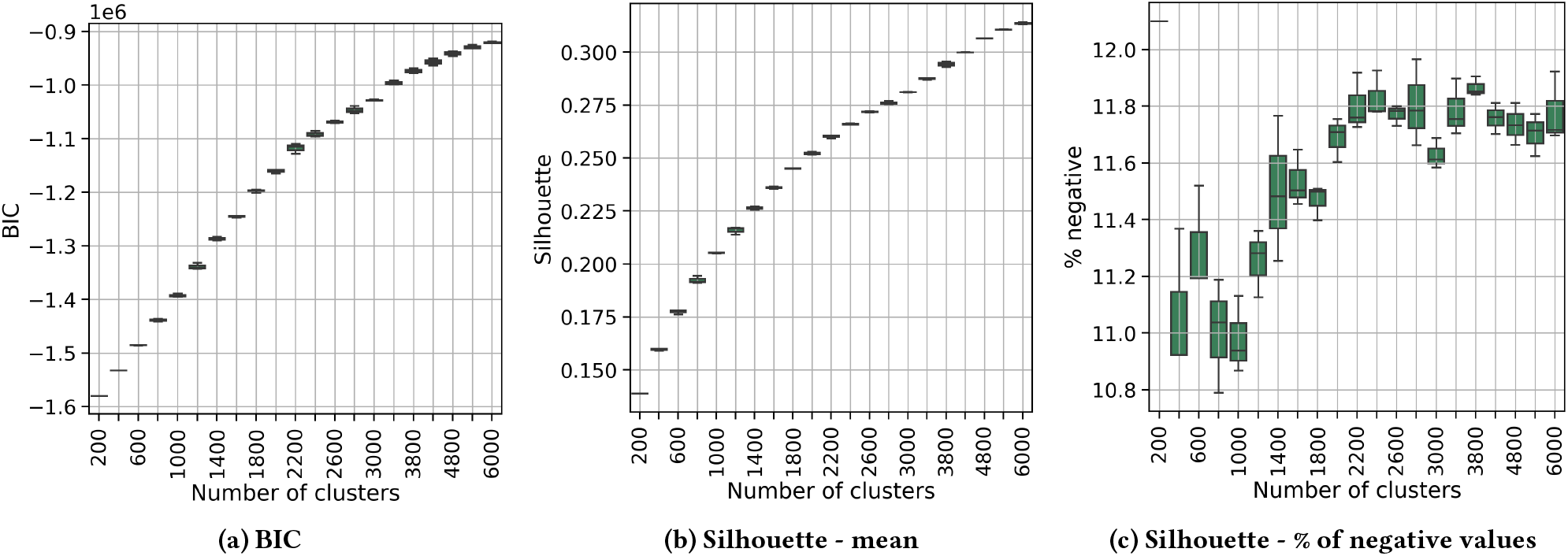
**Variation of BIC, mean silhouette scores and percentage of negative silhouette scores with respect to the number of clusters for *E. coli* for *k* = 6.**

Next, we evaluated the performance of OMclust as the value of *m* is varied, i.e., *m* = 1, .., 15. Figure 4 illustrates the precision, recall and *F*_*β*_ of this experiment. For lower values of *m*, the recall is higher since a large number of Rmaps are deemed to be related; however, this also results in lower precision. As *m* is increased, we find that the precision rises sharply while the recall drops. For both *k* = 4 and *k* = 5, we find the peak of the *F*_*β*_ curve is at *m* = 4, whereas, it is at *m* = 3 for *k* = 6. Based on this experiment, we choose *m* = 4 and *k* = 4 as the default setting for OMclust.

**Figure 4:**
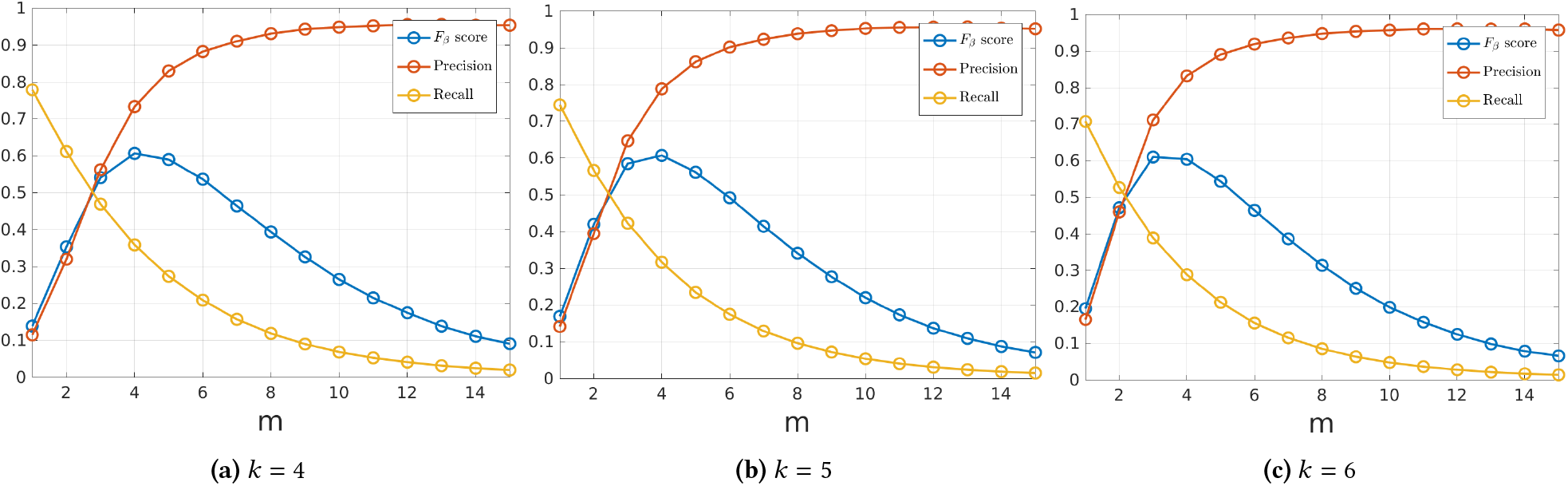
**Experimental results for simulated *E. coli* Rmaps using OMclust with varying values of *m* and *k*.**

As previously discussed, both Elmeri and cOMet use *m* and *k* in a synonymous manner as OMclust and therefore, we evaluated the performance of these methods as the value of *m* and *k* are varied. We note that the default setting is *k* = 5 and *m* = 10 for Elmeri, and *k* = 4 and *m* = 1 for cOMet. Figures 5 and 6 show the variation of precision, recall and *F*_*β*_ as these parameters are varied. Comparing, these figures with Figure Figure 4, we see that the maximum precision achieved by Elmeri is 50% and by cOMet is just under 60%; whereas the maximum precision achieved by OMclust is more than 90%. Moreover, recall of cOMet is consistently low (below 20%) when the precision is above 40%.

**Figure 5:**
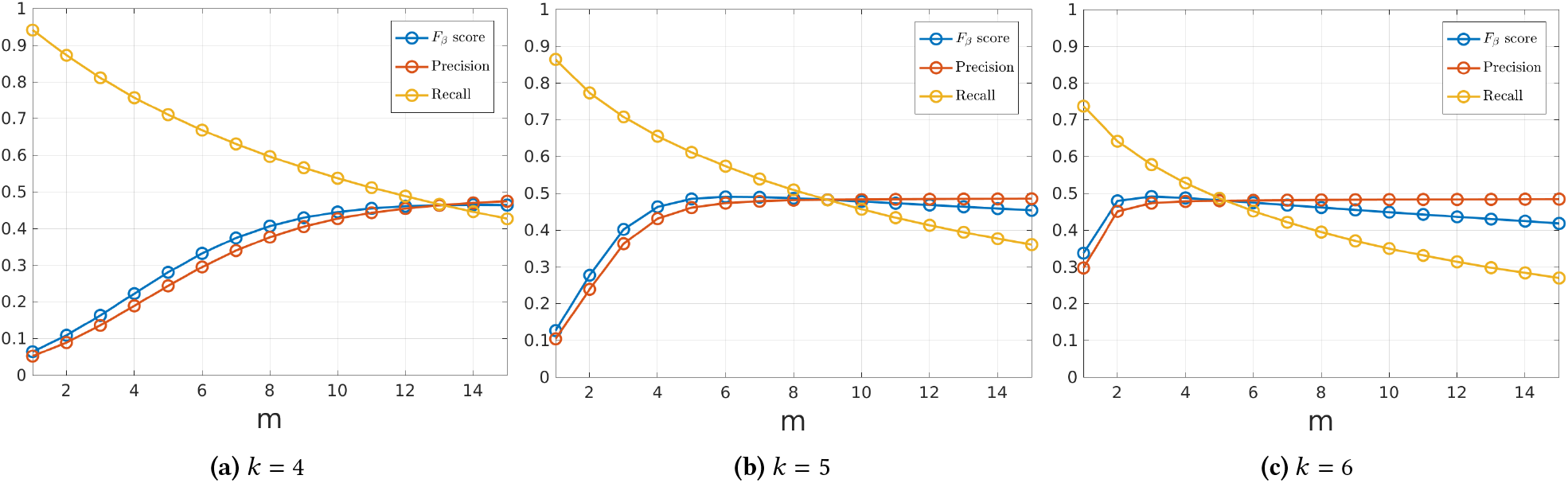
**Experimental results for simulated *E. coli* Rmaps of the heuristic for finding related Rmaps used by Elmeri, where *m* and *k* are varied**.

**Figure 6:**
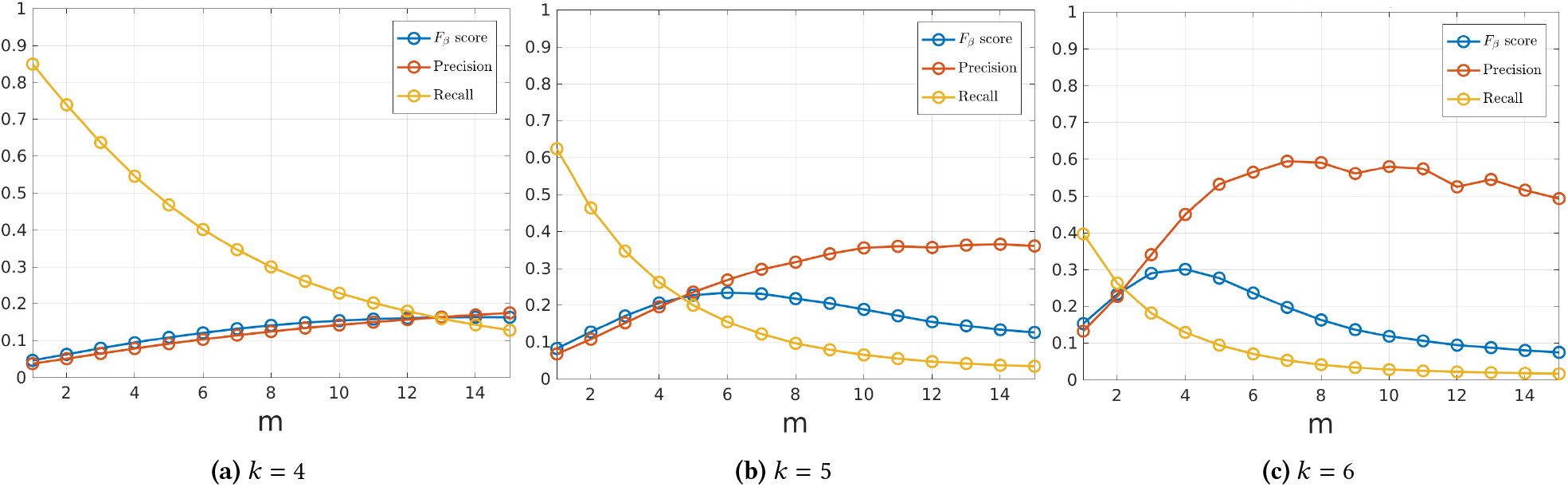
**Experimental results for simulated *E. coli* Rmaps of the heuristics for finding related Rmaps used by cOMet, where *m* and *k* are varied**.

Although we have shown that the BIC index should be robust (through the bootstrapping and the correlation with silhouette) to identify the optimal number range, the number of classes has an influence on the precision/recall. A lower number of classes improves recall at the price of precision. Since we chose the minimum number of clusters among those with a maximised BIC or within the region of stabilization after the elbow, the reported precision was the most conservative. Thus, if the cluster number was underestimated, the precision would only increase by choosing a higher number. The overestimation could still be an issue, but less probable, since in most datasets and for most values of *k*, there was a clear flexion of the BIC/silhouette.

In Table 1, we compare OMclust with cOMet and Elmeri using the *E. coli* test dataset. OMclust was ran with *k* = 4 and *m* = 4, and Elmeri and cOMet were ran with their default settings. OMclust required 6 seconds of CPU time and 0.01 GB of peak memory. cOMet had a CPU time of 10 seconds and peak memory usage of 0.74 GB, and Elmeri needed 5 minutes 49 seconds of CPU time and peak memory of 1.02 GB. OMclust had the highest precision (73.3%), followed by Elmeri with a precision of 48.3%, and cOMet with a precision of 5.1%. cOMet had highest recall (81.4%), followed by Elmeri (45.6%), and then OMclust (35.8%). Hence, cOMet had the highest recall but very low precision. OMclust had the highest precision but the lowest recall.

**Table 1:** **Comparison of the accuracy and efficiency of finding related Rmaps in simulated *E. coli* Rmap data. The peak memory (“peak mem”) is given in gigabytes (GB). The CPU time is reported in hh:mm:ss. OMclust was run with *k* = 4 and *m* = 4 and 2,200 clusters. Elmeri and cOMet were run with their default parameters, i.e., *k*=5 and *m* = 10 for Elmeri and *k* = 4 and *m* = 1 for cOMet.**

| Method | CPU time | Peak mem | Precision | Recall | No. of Related Rmaps |
| --- | --- | --- | --- | --- | --- |
| cOMET | 00:00:10 | 0.74 | 5.1% | 81.4% | 20,974,053 |
| Elmeri | 00:05:49 | 1.02 | 48.3% | 45.6% | 866,164 |
| OMCLUST | 00:00:06 | 0.01 | 73.3% | 35.8% | 445,121 |

Next, we compared the pairwise alignments of pairs of Rmaps that were predicted to be related from each of the methods. As previously mentioned, the S-score is the standard metric used to evaluate the alignment of two Rmaps. We note that higher S-score values represent larger and more consistent alignments. Therefore, we randomly sampled 100,000 pairs of Rmaps that were deemed to be related from each method and computed their pairwise alignments using the dynamic programming method of Valouev *et al*. Figure 7 illustrates the distribution of these values for the various methods. The mean S-score for cOMet, Elmeri and OMclust is 22.42, 33.52 and 35.01 with standard deviations 8.88, 12.20 and 14.55, respectively. Comparing the distributions and means, we see OMclust have a greater overlap compared to those found by the competing methods.

**Figure 7:**
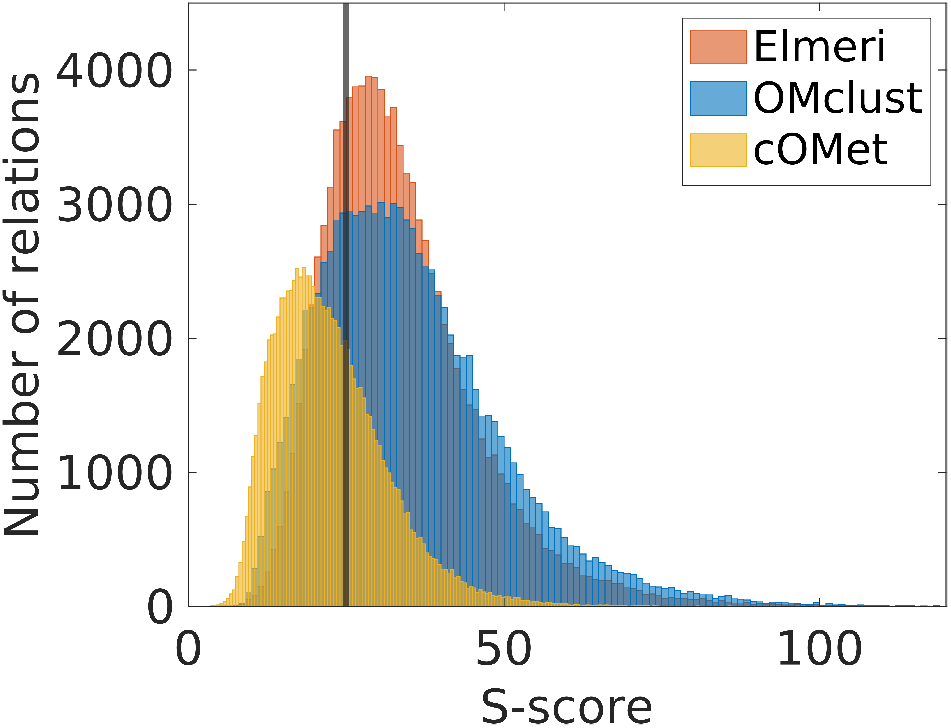
Comparison of the distribution of the S-scores of the related Rmaps found by Elmeri, cOMet and OMclust using simulated *E. coli* Rmap data. From each set of related Rmaps, 100,000 pairs of Rmaps were selected at random without replacement and pairwise aligned using Valouev *et al*. to calculate the S-score.

Further, we investigated the impact of using related Rmaps found by OMclust on error correction by integrating OMclust into cOMet and Elmeri. Table 2 shows the results of cOMet using the set of related Rmaps found by OMclust with various values of *m*. In addition, we show the performance of cOMet with its default setting. From these results, we conclude that our choice of *k* = 4 is confirmed since with this setting the best trade-off between CPU time, peak memory, and error correction performance is achieved. In particular, with *m* = 4, error correction improved the S-score of 5,303 Rmaps, compared to 5,301 for the default settings of cOMet; and the CPU time was reduced from 23 minutes 18 seconds to 16 minutes and 9 seconds. The mean S-score after error correction is correspondingly increased 67.66 to 68.03. We note that cOMet uses a heuristic to filter pairs of Rmaps that are unlikely to be related, which is not used when OMclust is used in combination with cOMet. Hence, although cOMet finds a larger number of relations, compared to what OMclust finds with *m* = 1 and *m* = 2, the CPU time of cOMet is slightly shorter.

**Table 2:** **Performance of cOMet using the related Rmaps found by OMclust with various values of *m*, i.e., *m* = 1, …, 10. In addition, we included the performance of cOMet using the default setting in the last row. The CPU time is in hh:mm:ss and peak memory (“Peak mem”) is in GB. “No. of improved Rmaps” is the number of Rmaps whose S-score increased after error correction. The mean and standard deviation of the S-score after error correction is also shown. We note the mean and standard deviation of the S-score for uncorrected Rmaps are 58.17 and 46.50, respectively**.

| m | No. of related Rmaps | CPU time | Peak mem | No. of improved Rmaps | Mean S-score | Std-dev of S-score |
| --- | --- | --- | --- | --- | --- | --- |
| 1 | 6,299,757 | 01:31:01 | 0.23 | 5,806 | 67.46 | 54.94 |
| 2 | 1,780,207 | 00:36:51 | 0.18 | 5,964 | 68.43 | 55.40 |
| 3 | 775,699 | 00:20:36 | 0.16 | 5,688 | 68.45 | 55.89 |
| 4 | 445,121 | 00:16:09 | 0.15 | 5,303 | 68.03 | 56.15 |
| 5 | 288,649 | 00:13:47 | 0.13 | 4,804 | 67.46 | 56.54 |
| 6 | 196,413 | 00:11:42 | 0.12 | 4,281 | 66.85 | 56.80 |
| 7 | 134,943 | 00:10:15 | 0.10 | 3,751 | 66.10 | 57.00 |
| 8 | 92,755 | 00:08:33 | 0.09 | 3,301 | 65.30 | 56.88 |
| 9 | 63,487 | 00:07:12 | 0.07 | 2,821 | 64.52 | 56.66 |
| 10 | 43,251 | 00:05:52 | 0.06 | 2,373 | 63.67 | 56.17 |
| COMET | 20,974,053 | 00:23:18 | 0.14 | 5,301 | 67.66 | 54.83 |

Similarly, Table 3 shows the results of Elmeri using the set of related Rmaps found by OMclust with various values of *m*. Here, we see that when OMclust is used in combination with Elmeri, it is able to obtain nearly the same mean S-score (70.2 verses 70.44) but in less than one third the CPU time and less peak memory.

**Table 3:** **Comparison of the performance of Elmeri using the related Rmaps found by OMclust using various values of *m*, and Elmeri with default setting (last row). The column headers are described in the caption of Table 2. The mean and standard deviation of the S-score for uncorrected Rmaps are 58.17 and 46.50, respectively**.

| m | No. of related Rmaps | CPU time | Peak mem | No. of improved Rmaps | Mean S-score | Std-dev of S-score |
| --- | --- | --- | --- | --- | --- | --- |
| 1 | 6,299,757 | 00:04:10 | 0.17 | 3,253 | 58.25 | 40.48 |
| 2 | 1,780,207 | 00:03:44 | 0.09 | 4,870 | 66.61 | 49.56 |
| 3 | 775,699 | 00:03:30 | 0.07 | 5,120 | 69.01 | 54.34 |
| 4 | 445,121 | 00:03:14 | 0.07 | 4,986 | 70.20 | 57.64 |
| 5 | 288,649 | 00:03:01 | 0.07 | 4,493 | 69.08 | 58.01 |
| 6 | 196,413 | 00:02:29 | 0.06 | 4,080 | 68.65 | 58.77 |
| 7 | 134,943 | 00:02:11 | 0.06 | 3,656 | 67.64 | 58.46 |
| 8 | 92,755 | 00:02:07 | 0.06 | 3,182 | 66.66 | 58.27 |
| 9 | 63,487 | 00:01:41 | 0.06 | 2,715 | 65.69 | 58.24 |
| 10 | 43,251 | 00:01:23 | 0.06 | 2,264 | 64.16 | 56.74 |
| Elmeri | 866,164 | 00:10:45 | 0.98 | 5,219 | 70.44 | 56.91 |

### 4.4 Results for Real Human Data

For training OMclust on the real human dataset, we simulated a dataset using the same restriction enzyme as used to generate the real data. We found the optimal number of clusters by performing a grid search from 10,000 clusters to 1,000,000 clusters, and plotting the BIC statistic, mean silhouette values, and percentage of negative silhouette values. The resulting plots are shown in Figure 8. Since the BIC and the mean silhouette do not have an inflection point but the percentage of negative silhouette values reaches a minimum at 100,000 clusters, we used this clustering. Training the model required 36,922 hours 31 minutes and 13 seconds of CPU time and 605 hours 5 minutes and 19 seconds of wall time running on 64 CPUs in parallel. The peak memory usage was 7.12 GB.

**Figure 8:**
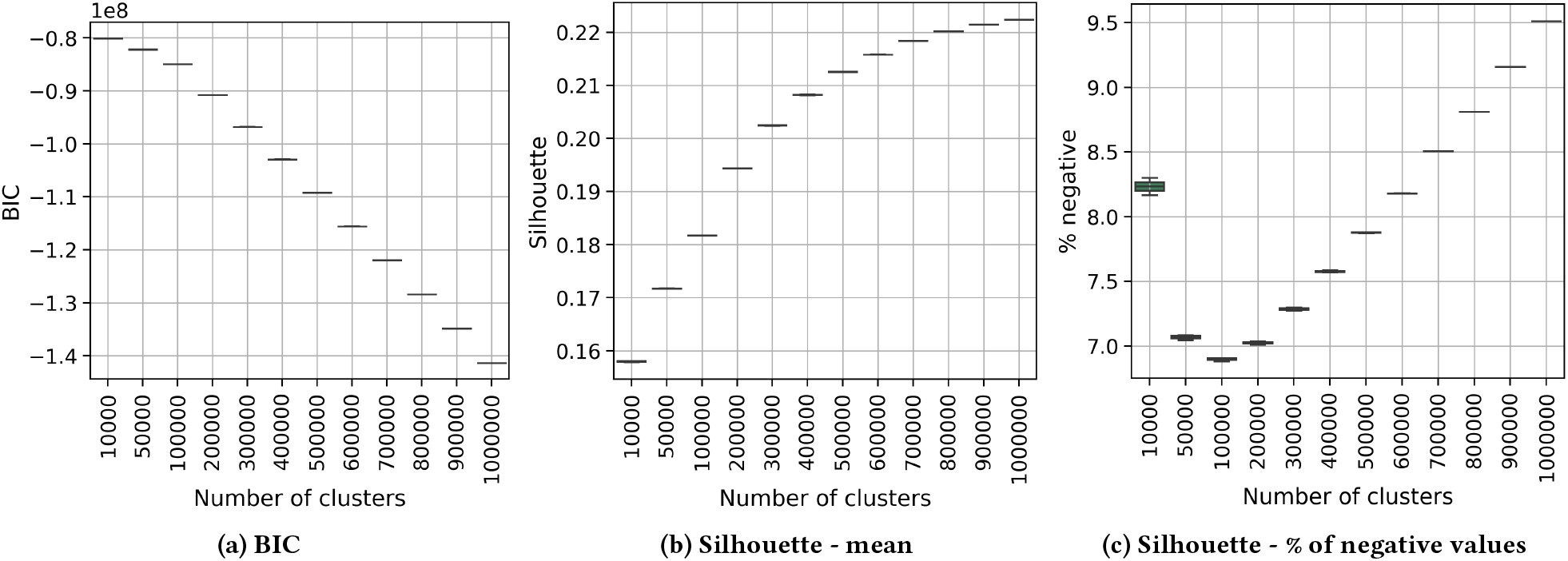
**Variation of BIC, mean silhouette scores and percentage of negative silhouette scores with respect to the number of clusters for human data**.

We evaluated our trained model using the real human data.

It took 2 hours 59 minutes and 17 seconds of CPU time to classify the *k*-mers into clusters with a peak memory of 10.48 GB and another 227 hours and 13 minutes and 44 seconds of CPU time to find related Rmaps. Because of the large number of Rmaps, we found related Rmaps on 1,000 CPUs in parallel which brought the wall-time down to just 22 minutes and 33 seconds with peak memory of 16.29 GB and found a total of 171,307,975 related Rmaps. For the classification step, we split the dataset into 1 ≤*X* ≤*N* streams where *N* is the total number of the Rmaps. For the *i*-th stream, we process *N* /*X* number of Rmaps, from e.g. from index *N* /*X*\* (*i* −1) to *N* /*X* \**i*. Each stream extracts the k-mers for each Rmap in its Rmap interval. The k-d-tree data structure used for classification is replicated in each stream, and used to assign each k-mer to a cluster center. All k-mer to cluster assignments are collected and distributed to all the streams. Each stream computes the *B* array from Section 3.3 which maps clusters to k-mers, and uses it to find the Rmaps related to the Rmap in its Rmap interval. The related Rmaps from each stream are written on a separate output file and these files are concatenated to produce the final output. Both Elmeri and cOMet were unable to run to completion on this dataset. Elmeri exceeded our memory constraint of 800 GB, and cOMet exceeded our disk space constraint of 10 TB. Elmeri crashed without any output; whereas, cOMet outputted the related Rmap pairs prior to crashing. Therefore, using this intermediate result of cOMet we sampled 100,000 pairs of Rmaps randomly without replacement from the pairs of Rmaps predicted to be related from OMclust, and from cOMet and calculated the S-score of these alignments using the method of Valouev *et al*. Figure 9 illustrates the distributions of these S-scores. The mean and standard deviation of the S-score of OMclust is 26.20 and 11.72, respectively; whereas for cOMet it is 18.21 and 8.74, respectively. We note that these distribution of cOMet does not reflect the distribution of a random sampling of all the relations and thus, may differ from the reported distribution – however, this would be unlikely as it would require a skewed distribution of the input data.

**Figure 9:**
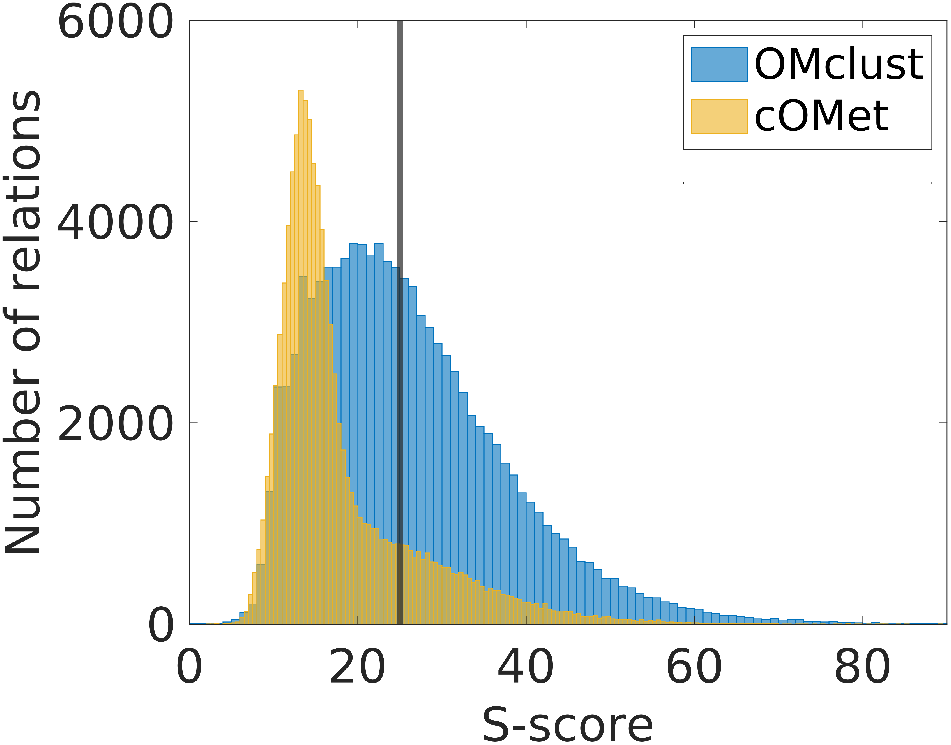
**Comparison of alignment scores of related Rmaps found by OMclust and cOMet for real human Rmaps. From the related Rmaps called by each method 100,000 relations were randomly sampled and the aligner of of Valouev *et al*. was used to align pairs of Rmaps from those relations. The alignment scores (called S-scores) were plotted as a histogram. The mean S-score for cOMet, and OMclust are 18.21 and 26.20 with standard deviations 8.74 and 11.72, respectively**.

Lastly, we compared the performance of error correction of cOMet by running it with its default setting and running cOMet using the set of related Rmaps found by OMclust. We note that although the initial step of finding all related Rmaps in cOMet ran out of disk space (see above), the error correction step of cOMet can still be ran since it does not use any external memory for this step. Both methods were run in parallel on 3,000 CPUs. As for the classification step, we split up the dataset into *X* streams and for each Rmap in the stream Rmap interval, we find related Rmaps, we align the Rmap to its related Rmaps and lastly, we error-correct the Rmap based on a consensus from the alignments. The error-corrected Rmaps from each stream are written on a separate output file and these files are concatenated to produce the final output. Each stream is handled as a separate job by an individual machine core. Table 4 shows the results of this experiment. We witness that the combined approach of using OMclust with cOMet uses a fraction of the CPU time and improves a significantly larger number of Rmaps. The mean S-score after error-correction improves from 37.41 to 59.01 using default cOMet, whereas it improves to 71.96 using OMclust relations. This demonstrates the benefit of using related Rmaps with high precision.

**Table 4:** **Error correction performance of cOMet on real human Rmap data using its default relations and using relations found by OMclust with *m* = 4 (OMclust + cOMet). The column headers are described in the caption of Table 2**.

| Method | Wall time | CPU time | Peak mem | No. of improved Rmaps | Mean S-score | Std-dev of S-score |
| --- | --- | --- | --- | --- | --- | --- |
| OMCLUST + cOMET | 00:50:25 | 638:41:35 | 16.29 | 2,757,266 | 71.96 | 44.87 |
| cOMET | 23:60:36 | 23637:21:11 | 15.73 | 987,985 | 59.01 | 25.18 |

## 5 CONCLUSION

In this paper, we developed a method for finding overlapping Rmaps that uses a Gaussian mixture model clustering, and does not require any quantization. We demonstrated that OMclust achieved the highest precision and was more efficient than competing methods, i.e., Elmeri and cOMet. In addition, we showed that using OMclust to find related Rmaps, the quality of error correction of cOMet is substantially improved and the CPU time is reduced. For example, the mean S-score of Rmaps generated from human DNA that were error corrected using cOMet with the related Rmaps from OMclust was 71.96; whereas the mean S-score was 59.01 using the default setting of cOMet. Moreover, the number of error corrected Rmaps also increased nearly 3x from 987,985 to 2,757,266 using OMclust. In summary, the high precision and efficiency of OMclust makes it suitable as a filtering step for finding related Rmaps for error correction, assembly or other applications of optical mapping data. We believe the integration of OMclust into these methods warrants future study. Another area that we warrants investigation is developing methods to select the optimal number of clusters. We used standard machine learning methods for cluster evaluation, which affects the time required for training the model. There may be a relationship between the coverage of the optical mapping data and the number of clusters, however, this relationship is complicated based on other factors, including the restriction enzyme and the repetition in the genome. However, predicting the number of clusters in a deterministic manner could potentially decrease the training time and thus, would be a valuable contribution.

## ACKNOWLEDGMENTS

This work was supported in part by the NSF IIS (grant number 1618814) and NSF SCH (grant number 2013998). Daniel Dole-Muinos and Ayomide Ajayi were supported by the NSF REU program.

